# GraphOmics: An Interactive Platform To Explore And Integrate Multi-Omics Data

**DOI:** 10.1101/2021.06.24.449741

**Authors:** Joe Wandy, Ronan Daly

## Abstract

**Background:** An increasing number of studies now produce multiple omics measurements that require using sophisticated computational methods for analysis. While each omics data can be examined separately, jointly integrating multiple omics data allows for a deeper understanding and insights to be gained from the study. In particular data integration can be performed horizontally, where biological entities from multiple omics measurements are mapped to common reactions and pathways. However, data integration remains a challenge due to the complexity of the data and the difficulty in interpreting analysis results.

**Results:** Here we present GraphOmics, a user-friendly platform to explore, integrate multiple omics datasets and support hypothesis generations. Users can upload transcriptomics, proteomics and metabolomics data to GraphOmics. Relevant entities are connected based on their biochemical relationships, and mapped to reactions and pathways from Reactome. From the Data Browser in GraphOmics, mapped entities and pathways can be ranked, sorted and filtered according to their statistical significance (p-values) and fold changes. Context-sensitive panels provide information on the currently selected entities, while interactive heatmaps and clustering functionalities are also available. As a case study, we demonstrated how GraphOmics was used to interactively explore multi-omics data and support hypothesis generations using two complex datasets from existing Zebrafish regeneration and Covid-19 human studies.

**Conclusions:** GraphOmics is fully open-sourced and freely accessible from https://graphomics.glasgowcompbio.org/. It can be used to integrate multiple omics data horizontally by mapping entities across omics to reactions and pathways. Our demonstration showed that using interactive explorations from GraphOmics, interesting insights and biological hypotheses could be rapidly revealed.

## Background

The availability of high-throughput technologies means many studies are increasingly producing large-scale untargeted measurements of different biological entities, such as transcripts, proteins and metabolites. Combining the diverse set of omics data produced from different measurement platforms is often required as the initial step of an integrated analysis. Data integration has been shown to reveal stronger findings compared to analysing a single dataset alone, with wide-ranging successes from studying the human microbiome to identifying cancer biomarkers Misra et al. [2019], Lloyd-Price et al. [2019], Vasaikar et al. [2018].

Omics integration approaches can be divided into two types: vertically where integration is performed by using multiple omics data from the same biological sample; and horizontally where the integration is performed by mapping shared or related entities from different biological samples Jendoubi [2021]. One popular approach to vertical integration is through matrix factorisation. This includes methods such as Canonical Correlation Analysis (CCA) that finds canonical variables maximally correlated to each other from the different omics data, as well as data fusion via tri-matrix factorisation Žitnik and Zupan [2014] that considers the relations and constraints across and within omics, and decomposes the data into low-rank matrices that reveal hidden associations. Another example is Multi-Omics Factor Analysis (MOFA) Argelaguet et al. [2018] that provides a Bayesian model and a robust inference scheme to factorise omics data into latent factors explaining the main variations in the data.

During vertical integration, often it is required for different omics measurements from the same sample to be matched. However in some instances, existing data cannot be matched in this manner, since not all omics types were measured or due to other limitations in the study. Horizontal integration offers an alternative scheme, where integration is performed by mapping shared or related entities from one omic dataset to another without requiring for samples to be aligned. Instead biological pathways could serve as the shared context onto which entities are mapped.

In recent years, Web-based tools to perform horizontal integration using pathways have been gaining popularity. For example MetaboAnalyst Pang et al. [2021], considered one of the most popular online tools in metabolomics at the time of writing, provides a functionality to map genes and metabolites to metabolic pathways and performs pathway enrichment analysis. Another example is 3Omics Kuo et al. [2013] which accepts human-only transcriptomics, proteomics and metabolomics datasets and perform pathways mapping as well as other analyses such as correlation and gene ontology (GO) analyses. Finally PaintOmics3 Hernández-de Diego et al. [2018] performs a complete integration of multiple data types to KEGG pathways, allowing for the enrichment and clustering analyses of pathways, as well as network visualisation.

Despite this abundance of tools, data integration remains a challenge due to the complexity of the data, and the difficulty in relating analysis results to biological interpretations. A common approach employed by many tools is to present analysis outcome as a complex network graph Pang et al. [2021], Kuo et al. [2013], Hernández-de Diego et al. [2018], Cottret et al. [2010]. Networks are visually appealing as unstructured results can be easily rendered as a graph having nodes and edges. Nodes represent different biological entities, while the relationships between nodes can be flexibly represented by edges that capture different interactions between the nodes. However the complexity of a typical multi-omics study means networks can quickly grow to a large size, having numerous nodes and edges. When biologists are presented with a ‘hairball’ network, deciphering biological meaning and generating hypothesis from such outputs can be challenging Schulz and Hurter [2013]. A similar challenge is also faced in interpreting analysis results presented as long and static (non-interactive) tables.

Here we introduce GraphOmics, a Web application that accepts measurements of transcripts, proteins and metabolites and perform data integration horizontally using Reactome Croft et al. [2014] as the graph knowledge base. GraphOmics provides an interactive platform that integrates data to Reactome pathways emphasising interactivity and biological contexts. This avoids the presentation of the integrated omics data as a large network graph or as numerous static tables. Instead each biological entity is mapped onto Reactome reactions and pathways using biochemical knowledge, and presented in the context of their relationships to other related entities. Interactive explorations of linked entities form the centrepiece of GraphOmics, where selecting an entity will display other entities related to it. Further analyses such as gene ontology enrichment and pathway analysis spanning multiple -omics data can be performed. Finally biological conclusions can be annotated in GraphOmics and the results shared to others.

## Implementation

Figure 1 provides a diagram of overall GraphOmics functionalities. An initial data loading step is performed to get measurements of entities into GraphOmics. As part of data loading, the Reactome database is used for mapping of the biological entities (transcripts, proteins and metabolites) in the uploaded data onto reactions and pathways from Reactome. Once data loading is completed, users can perform various global analyses, including differential analysis, pathway activity enrichment, principal component analysis (PCA), clustering and uni-variate statistical tests for differential analysis. To assist in data interpretation, mapped results are shown in multiple interactive tables that are linked to each other. Selecting an entry in one table will filter entries in other related tables. Groups of related entities can also be created and analysed within GraphOmics.

**Figure 1:**
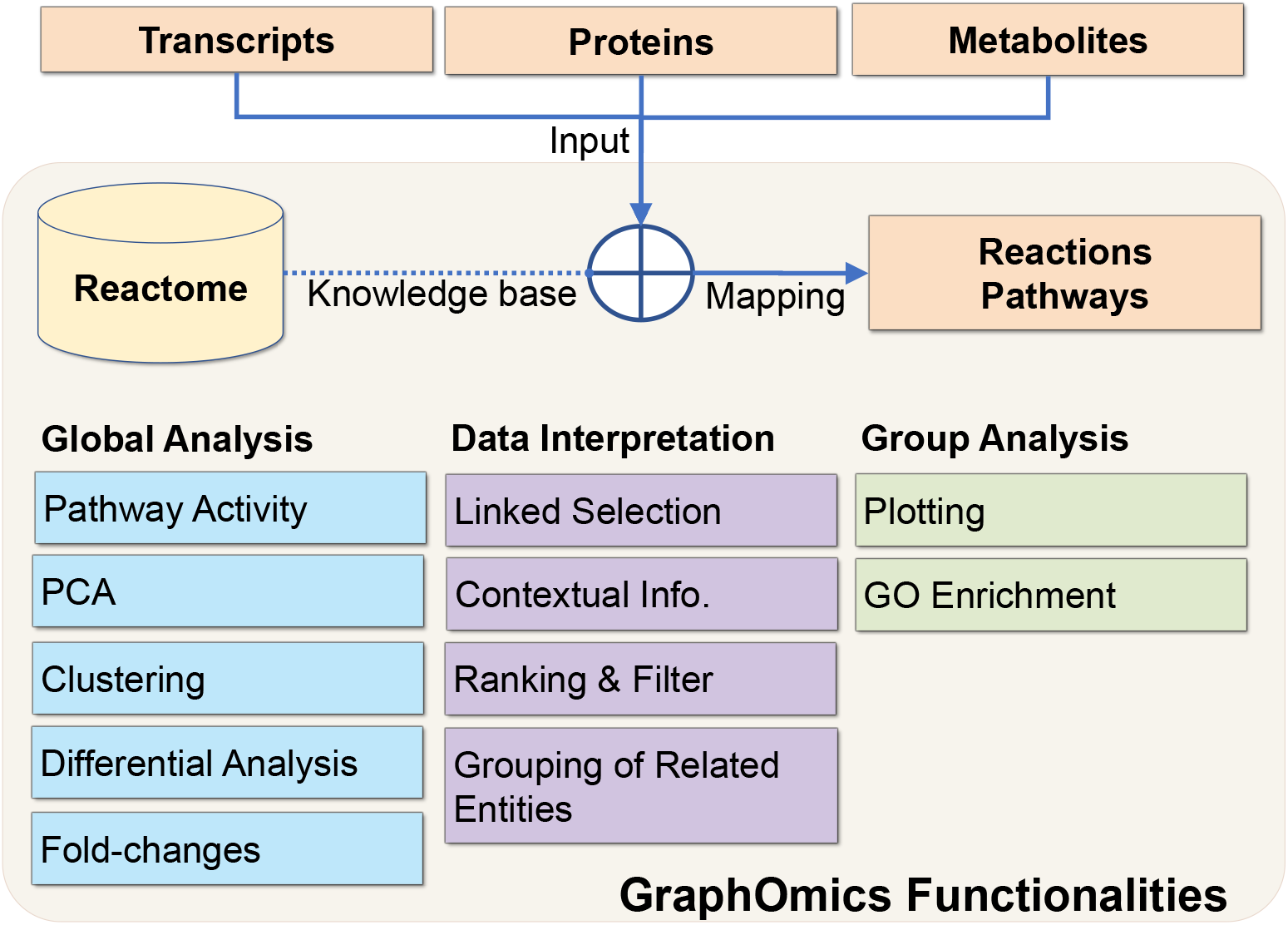
Overall GraphOmics functionalities. Horizontal integration is performed in GraphOmics by mapping transcripts, proteins and metabolites to Reactome’s reactions and pathways. From the platform, global analyses can be performed and data interpreted in in an interactive manner.

### Overall system design

GraphOmics is a Web-based system developed using open-source technologies. The client (browser) side is built upon HTML & Javascript, while charting functionalities are provided through libraries such as D3 and Plotly. The server side runs on Django 2 Web framework and the Python 3 programming language. Common statistical methods such as t-tests and PCA are implemented using the numpy and scipy libraries in Python, while differential analyses using *DeSEQ2* [Love et al., 2014] and *limma* [Smyth, 2005] are provided through R. An SQLite database is used to store relational data. A local copy of Reactome knowledge base [Croft et al., 2014] is downloaded and accessed from the Django Web application through a Neo4j graph database.

### Data loading and mapping

To begin analysis, users upload their transcripts, proteins or metabolites data to GraphOmics. Uploaded data are provided as matrices in a Comma-separated Value (CSV) format, where rows in the matrix are the entity IDs, columns are the samples, and entries are the measurements. Additionally users can also include the results for any statistical test that has been performed outside GraphOmics, in form of fold-changes and statistical significance (p-values) of entities.

To facilitate mapping, GraphOmics requires each row in the CSV file to be labelled with the appropriate ID for that entity type. For transcripts, the Ensembl ID of the gene that produces the transcript should be used. For protein data, the UniProt ID of the protein is required. Finally for compound data, the KEGG or ChEBI IDs of the compounds are accepted. Additionally users also provide the sample names and a design matrix specifying the assignment of samples to experimental conditions (refer to Supplementary S1 for details on the input format).

Horizontal integration of the uploaded data is performed through an automated mapping procedure written in Cypher (the graph query language used in Neo4j). This retrieves the connections between transcripts, proteins, metabolites to reactions and pathways of the given species in Reactome, constructing a network graph of entities, reactions and pathways involved in the dataset. Entities in this network graph are connected to one another: transcripts are linked to the proteins they encode, proteins and compounds are linked to the reactions they are involved in, and reactions are linked to the pathways that contain them. Once the initial mapping is completed, the results are stored in the SQLite database and presented to users in the Linked Data Browser.

### Multi-Omics Data Interpretation

#### Linked Data Browser

The Data Browser is the primary interface in GraphOmics that facilitates linked exploration of the integrated data. Instead of presenting an often-massive network graph, the main components of the Data Browser are five interactive tables: one for each supported omics type (transcripts, proteins and metabolites) as well as for reactions and pathways (Figure 2).

**Figure 2:**
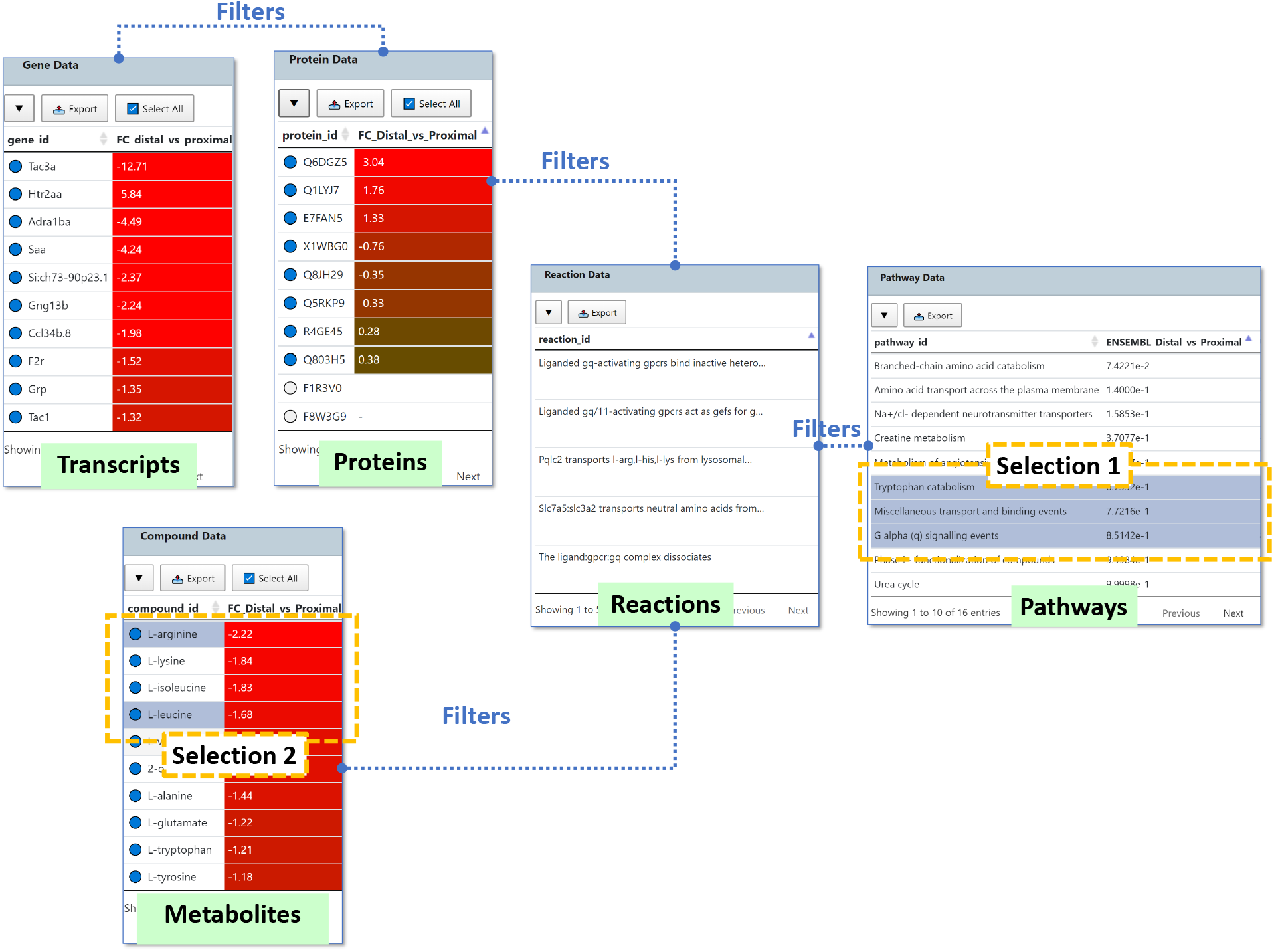
Data Browser in GraphOmics. The Data Browser in GraphOmics facilitates linked explorations of multi-omics data. Transcripts are linked to the proteins they synthesise. Proteins and metabolites are linked by reactions they are involved in. Reactions in turn are linked to Pathways that contain them. Entries in all tables can be selected by clicking on them. Selections are used to filter entries in other linked tables. Multiple tables can be selected in turn to define a flexible filtering criteria. For example, selecting the three pathways (Selection 1) will filter for reactions, proteins, metabolites and transcripts that are connected to the selected 3 pathways. If the user subsequently select two metabolites (Selection 2) from the filtered results, the results are further filtered to include only transcripts, proteins and reactions connected to the two selected metabolites under the 3 initially selected pathways. Each table can also be searched, sorted and filtered according to their fold changes and p-values. Blue circles next to the entity name indicate measured entities.

Users interact with the Data Browser by navigating through the tables. Clicking an entity in the Data Browser selects it, and multiple entities can be selected in this manner. Selections from one table will filter entries in other tables, such that only connected items are shown according to the links between entities. As more entities are added to the current selection, the number of entities displayed across tables are reduced to meet the filtering criteria.

In this manner, users can explore the data starting from a global view where all entities are shown, and successively narrowing down to more specific entities that are related to the selected items. This ‘drill-down’ interactivity in the Data Browser could help reveal the relationships among biological entities of interest and their reactions and pathways across omics.

#### Contextual Information Panel

Selected entries in the Data Browser are also associated to contextual information under each table (Figure 3). This includes plots of the measurements of that entity across conditions as well as links to external databases (Figure 3A,B). For transcripts, the Harmonizone Web service [Rouillard et al., 2016] is used to retrieve additional description for the gene, as well as links to Ensembl and GeneCard. For proteins, the name, catalytic activity, pathways, gene ontology terms, and links to Uniprot and Swiss-Model of the currently selected proteins are displayed. For compounds, information on the KEGG and CheBI IDs, formula and SMILES string, as well as links to their respective databases, and also compound structures are retrieved. For reactions and pathways, a desriptive summary is displayed by querying Reactome (Figure 3C). Additionally an interactive pathway viewer utilising the Reactome Pathway Diagram Viewer (DiagramJS) is also available (Figure 3D). Measured values of transcripts, proteins and metabolites can be overlaid on top of the interactive pathway diagrams.

**Figure 3:**
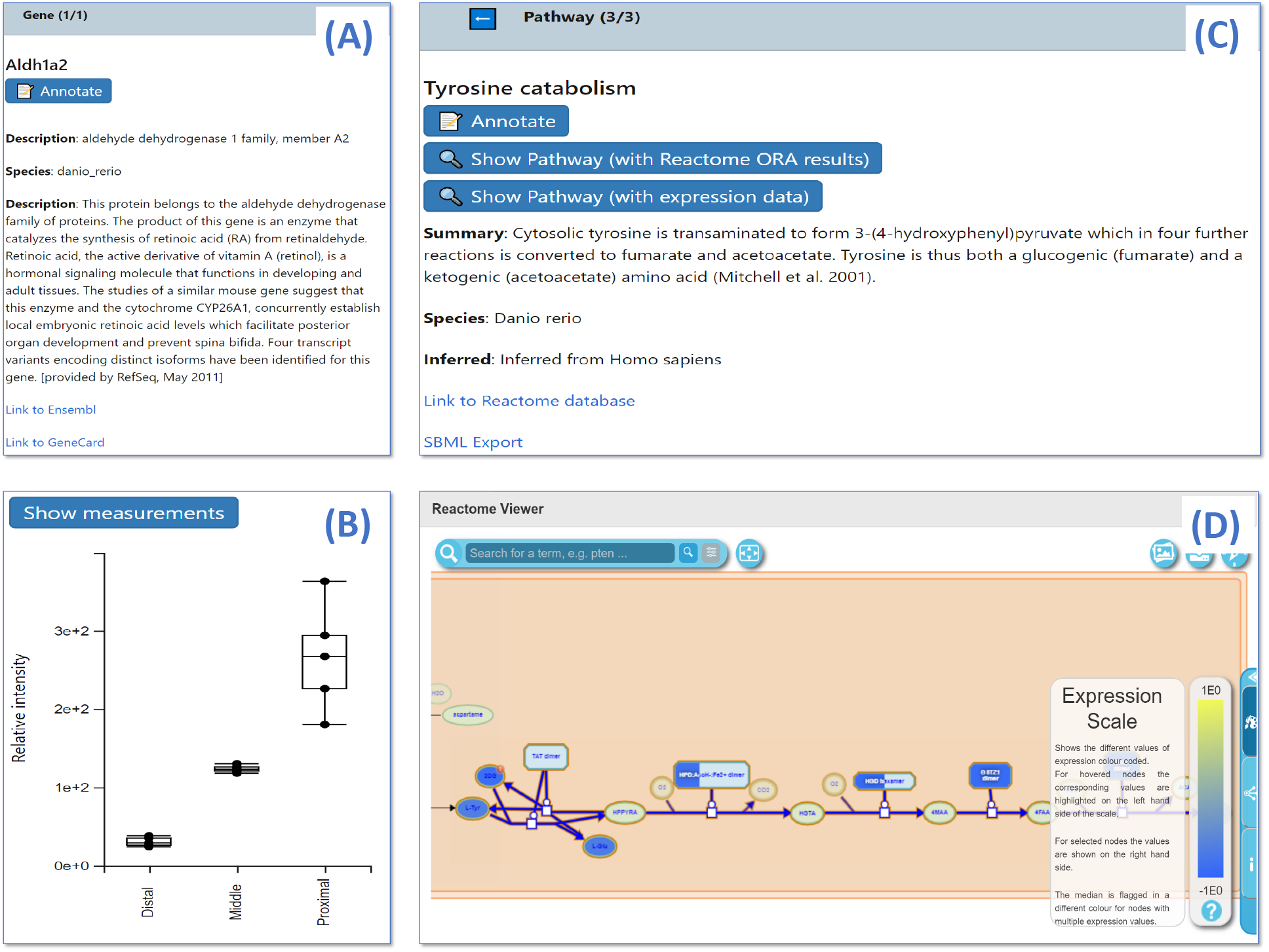
The Info Panel in GraphOmics. The info panel provides additional contextual information for selected entries in the Data Browser. **(A)** An example info panel entry for the transcript identified by the gene *Aldh1a2*, as well as **(B)** its measurements if available. Entities and pathways can be annotated by clicking on the *Annotate* button in the Info Panel. **(C)** An example info panel entry for the *Tyrosine catabolism* pathway. Clicking the *Show Pathway* button displays **(D)** an interactive pathway diagram via DiagramJS, with either Reactome ORA results or expression data mapped onto it.

#### Ranking and Filtering

All interactive tables in the Data Browser allow entities to be ranked and sorted according to their fold changes and p-values. This can be used to explore the most significantly changing entities across omics that are differentially expressed (DE). In conjunction with linked interactions, the interface allows users to easily navigate through the top DE entities from one omics and inspect if they are linked to DE entities from other omics. Entities are also connected to pathways, which can be subjected to enrichment analysis within GraphOmics. In this manner, users can easily rank DE entities and determine which enriched pathways they are connected to. Additionally the Query Builder in GraphOmics allows for complex queries to be defined on the data (Figure 4). From the Query Builder, a query can be defined using comparison operators to filter entities by their p-values and fold changes. Queries spanning multiple omics data can also be defined by concatenating (performing a logical AND operation) of each constituent single-omics query.

**Figure 4:**
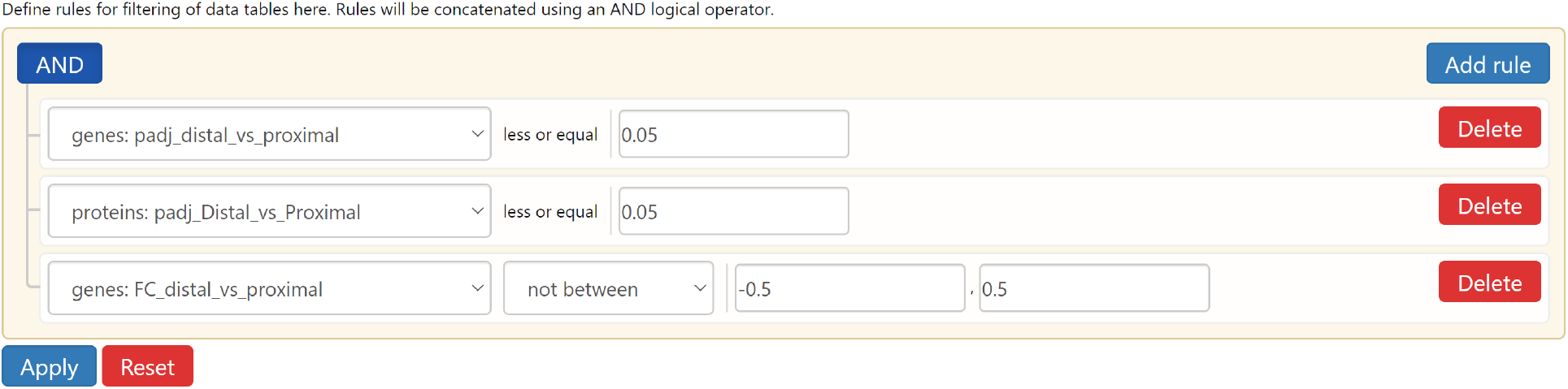
The Query Builder in GraphOmics. The Query Builder is used to filter entities of data tables by specifying rules that will be concatenated using a logical AND operator. In this example, a query is constructed to filter for transcripts and proteins that are both statistically significant (p-values less than 0.05) and having transcript fold changes at least 0.5 both ways.

#### Creating and Analysing Groups

GraphOmics allows for any set of entities that have been selected by users to be saved as a selection group. These groups can later be loaded for future use. A group of related entities (for instance the top DE entities, or members of a cluster or some pathways of interest) can be defined, saved and loaded for future analysis. Selection groups can be easily visualised and plotted. For transcriptomics data, gene ontology analysis can be performed using the Python package GOATools [Klopfenstein et al., 2018] to discover enriched GO terms associated with a group. Additionally interactive heatmaps and clustering analysis using Clustergrammer can also be performed on any group. Finally users can annotate groups on the GraphOmics platform for reporting purposes.

### Global Analysis of Multi-Omics Data

#### Differential Expression Analysis

A common task in omics data analysis is to find entities that are differentially expressed (DE) across different experimental conditions. If users have performed their own DE analysis, the statistical significance (p-values) of entities could be uploaded as part of data loading process. Otherwise from the Inference page in GraphOmics, users can execute standard uni-variate t-tests (with Benjamini–Hochberg procedure for controlling the false discovery rate). Additionally, widely-used methods such as *DeSEQ2* and *limma* can also be run as an option. The resulting statistical significance from performing DE analysis are shown in the interactive tables of the Data Browser, alongside the entity names and measured values.

### Interactive Clustering and Heatmap

Heatmap visualisation is performed using Clustergrammer [Fernandez et al., 2017], a Web component that integrates interactive heatmap and hierarchical clustering to visualise high-dimensional biological data. Clustergrammer provides many interactive features to explore a hierarchically clustered heatmap, including navigational features such as zooming and panning, as well as filtering features to search and select entities.

The interactivity of Clustergrammer makes it suitable for integration with GraphOmics as it works in concert with the Data Browser. Each omics type (transcripts, proteins and metabolites) in the Data Browser is associated to a Clustergrammer component (Figure 5). Clustergrammer was modified such that selecting entities in the Data Browser also performs the same selection in the corresponding Clustegrammer component, and vice versa.

**Figure 5:**
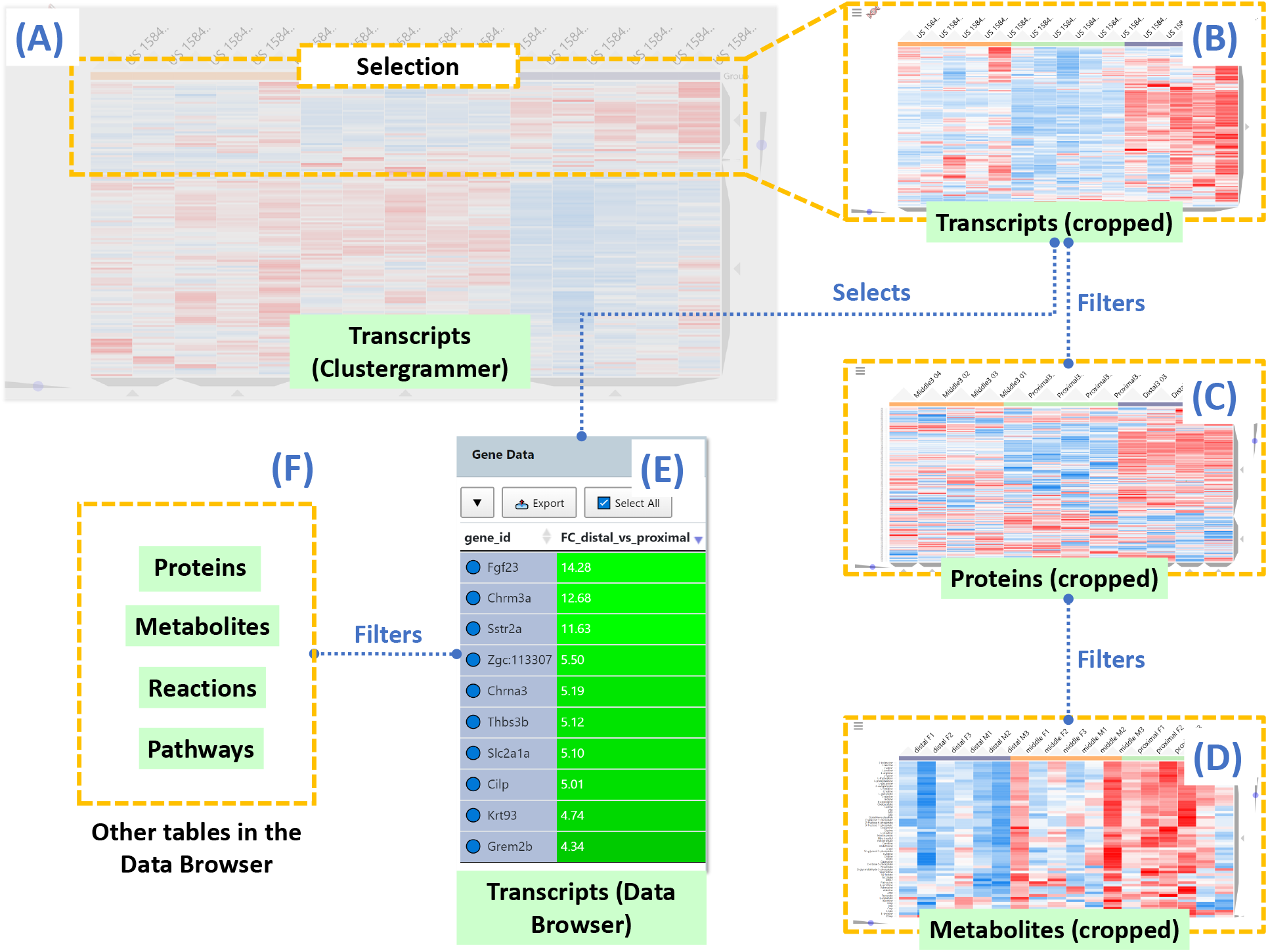
Clustergrammer Integration in GraphOmics. **(A)** Clustergrammer displays a hierarhically-clustered interactive heatmap, where clusters can be selected at any level of the dendogram. For example, here we show an example Clustergrammer component for the transcripts Zebrafish data. **(B)** Selecting a cluster in the Clustergrammer will display a cropped view of that data. For example, here we show an example cropped Clustergrammer showing only transcripts in the currently selected cluster. **(C, D)** Entities in related Clustergrammers are also filtered according to their relationships to the selected entities. **(E)** Entities in the selected cluster are also selected in the corresponding Data Browser table. **(F)** This in turn will filter other related tables in the Data Browser. The selection process can also be performed in reverse such that selecting entities in the Data Browser also filters the linked Clustergrammers (going backward from E to A in the diagram).

Clustergrammer integration means users can generate a heatmap and perform cluster analysis for any selections in the Data Browser. For instance, this includes the ability to display the heatmap of entities in a pathway (or in several pathways), or to discover the clusters of proteins and metabolites linked to top DE transcripts. The interaction also goes the other way, such that selecting a cluster in Clustergrammer also selects its member entities in the Data Browser. This allows users to examine the DE members of a cluster and their connections to reactions and pathways.

### Principal Component Analysis

PCA can be used to assess the global similarity of samples across different conditions. In GraphOmics, PCA analysis is created from the Inference page by selection the omics type and as the number of components to use. The results from PCA analysis include plots of the projected samples for the first two principal components, as well as a scree plot showing the percentage of variance explained by the different components. The latter plot can be examined to determine how many components to retain for analysis.

### Pathway Activity Analysis

Enrichment of a pathway often suggest relevant biochemical activities happening in that pathway. In GraphOmics, pathway activity analysis can be performed by considering a single omics dataset separately, or from multiple omics datasets at once. To prioritise changing pathways in single omics data, we developed a Python library named PALS [McLuskey et al., 2021] that presents a unified wrapper to the following algorithms: Over-representation Analysis (ORA); Gene Set Enrichment Analysis (GSEA) [Subramanian et al., 2005]; and Pathway Level Analysis of Gene Expression (PLAGE) [Tomfohr et al., 2005]. Originally developed for metabolomics, PALS was extended in GraphOmics to be able to deal with transcripts and proteins data too.

The three pathway ranking methods in PALS represents a diverse approach to enrichment analysis. ORA is widely used to assess the probability of over-representation of DE entities in a pathway using the Hypergeometric test. GSEA is considered a ‘second-generation’ method that takes into account the correlation between sets of entities to assess DE pathways. Finally PLAGE is a method based on singular value decomposition found to be best performing in [Tarca et al., 2013] returning the highest detection of changing pathways.

From the Inference page, users can choose to run any of these methods on the GraphOmics server. For any of the pathway ranking method, the p-values of significantly changing pathways are collected and displayed with pathway names in the Data Browser. This lets pathways to be ranked, sorted and filtered in the same manner as entities.

### Multi-Omics Pathway Activity Analysis

GraphOmics offers a way to perform pathway analysis separately on each omics, and integrate the results at the end. The separate pathway analysis results run on different omics datasets can be combined with an AND operator in the Query Builder. For instance from the Query Builder, users can easily filter pathways that are significantly changing based on the transcripts AND proteins AND metabolites measurements.

For a different approach that considers multiple omics data together during analysis, users can run the Reactome Analysis Service, which offers a high-performance multi-omics over-representation analysis using the Reactome server [Fabregat et al., 2017]. The IDs of DE entities (across multiple omics) are selected according to a user-defined threshold on the p-values, which defaults to ≤ 0.05. The collected IDs of DE entities is sent to Reactome Analysis Service, which performs pathway analysis through ORA on the Reactome server. Analysis token is returned, and the results of DE pathways and their p-values are retrieved in GraphOmics and displayed on the Data Browser for sorting and filtering.

## Results

### Comparison to Other Multi-Omics Systems

A comparison of GraphOmics to several other popular Web-based multi-omics systems, namely MetaboAnalyst [Pang et al., 2021], 3Omics [Kuo et al., 2013] and PaintOmics3 [Hernández-de Diego et al., 2018], is provided in Table 1.

**Table 1:**
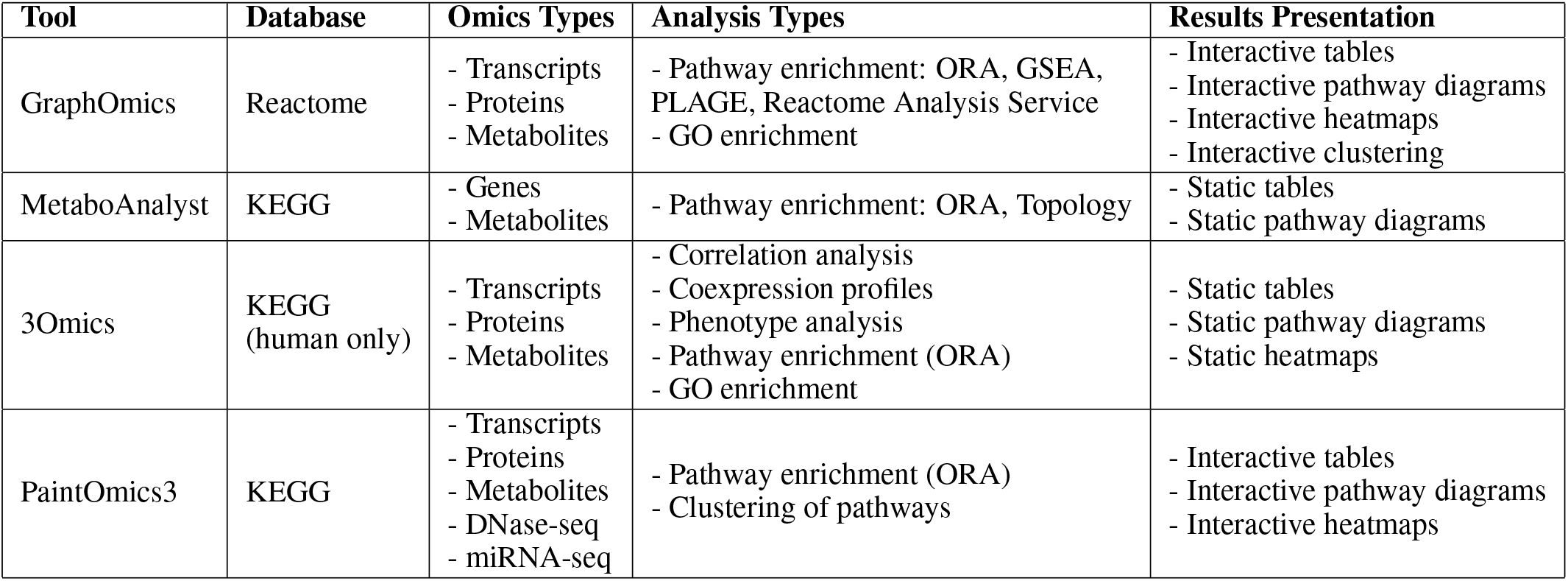
A comparison of GraphOmics to other Web-based multi-omics systems.

All systems evaluated provide a functionality to map a list of identifiers and associated measurements to pathways. GraphOmics relies on the Reactome database, while the others use KEGG. 3Omics is limited to the analysis of human data only, while the other systems evaluated, including GraphOmics, can handle many species. All systems provide a way to rank and prioritise relevant pathways using either single or multiple omics datasets. ORA appears to be the most common method for ranking pathways, although MetaboAnalyst provides an option that considers the topology of pathways during analysis. Additionally 3Omics provide many functionalities not directly related to pathways, such as correlation analysis, that could be useful in revealing interesting biological entities.

Both MetaboAnalyst and 3Omics generate analysis results as static tables and graphs. The large amount of non-interactive results produced by MetaboAnalyst and 3Omics could potentially be difficult for users to navigate. PaintOmics3 could be considered closest to GraphOmics in interactive functionalities. Analysis results are presented in PaintOmics3 as a sorted interactive table or as a network graph of pathways, with nodes representing significant pathways and edges drawn based on their linked biological processes. ‘Painting’ a pathway reveals additional information for that pathway, including the pathway diagram and an interactive heatmap showing measured values. PaintOmics3 also offers a novel analysis where pathways with similar trends can be clustered. Clustering results are overlaid on the network graph to reveals groups of pathways with similar changes.

GraphOmics differs in several key aspects when compared to PaintOmics3: our interface allows data explorations to begin from any entity of interests (for instance starting from the top DE transcripts), while in PaintOmics3 explorations are centered around DE pathways as the starting point. The linked views in GraphOmics reveal the explicit individual connections between all connected entities for easy inspections, while in PaintOmics3 these connections are summarised as edges between pathways in the network graph. From the Information Panel, GraphOmics displays more contextual information for each selected entity than PaintOmics3. Integration with Clustergrammer also means any clusters of entities can be identified and visualised as heatmaps, and their connections to others displayed in the Data Browser. This is a capability not present in PaintOmics3.

### Zebrafish Case Study

Using a public Zebrafish dataset [Rabinowitz et al., 2017], we demonstrated how biological insights could be gained through data integration and interactive explorations in GraphOmics. The aim of the original study was to uncover relevant biomarkers that regulate patterned regeneration in Zebrafish fins. This process is regulated by positional memory allowing cells to be regenerated at their previous locations before injury.

### Data Loading and Pre-processing

The processed transcripts, proteins and metabolites data from the original study was retrieved. For each omics type, a measurement CSV was created where rows corresponded to the entities and columns were the samples. Each row was identified by a unique identifier column, with ENSEMBL gene ID, UniProt ID and KEGG ID used for identifying transcripts, proteins and metabolites respectively. Positional memory is established by molecules that exist in a gradient along the uninjured appendages, so the measured samples were divided into three experimental conditions according the proximity in the fins where the sample was obtained: proximal, middle and distal (with proximal the closest to the torso and distal the furthest). Following the original study, we focused on the comparison of distal-vs-proximal where the largest differences could be seen.

CSV files for the multi-omics Zebrafish data was uploaded to GraphOmics. Automated mapping was performed by GraphOmics, resulting in 8690 transcripts linked to 8010 proteins and 462 compounds across 6995 reactions and 1272 Reactome pathways. The original processed transcriptomics data already contained DeSEQ2 analysis results comparing distal to proximal which were retained during upload and used as the DE results for the transcripts. This demonstrates how additional analysis from an external workflow could be easily incorporated into GraphOmics.

Differential expression analysis is often used to highlight significantly changing entities that could be of biological interests. From the original study, DE results were already available for the transcripts and so they were used. For the protein and metabolite data, we employed *limma* to perform the DE analyses of proteins and metabolites. PLAGE was used to perform DE analysis of pathways using each omics data separately as the input, resulting in different sets of p-values for each pathway depending on the source data used. This was all performed from the Inference tab in GraphOmics. All results from DE analysis in form of p-values and fold-changes (if available) are displayed in the Data Browser alongside the entities.

### Interactive Omics Exploration of the Zebrafish Data

Here we showed how GraphOmics easily characterised the set of DE transcripts linked to DE proteins. This could be used to identify the important transcripts and proteins that are involved in establishing positional memory of zebrafish. The following query was formulated from the Query Builder: filter for transcripts and proteins with a threshold of 0.05 on the p-values, and having at least ±0.5 on the log fold changes of the transcripts (Figure 4). The results were a selection of 87 transcripts and their corresponding proteins, as well as 21 compounds involved in reactions catalysed by those proteins. Note that the automatic mapping approach in GraphOmics revealed 11 out of the 32 DE transcripts linked to DE proteins in the original study in [Rabinowitz et al., 2017]. Among the DE transcripts found in agreement with the original study were the gene *aldh1a2* which catalyses the synthesis of retinoic acid, as well as *muc5.2* found to be retained in both uninjured and early stages of injuries. Both genes were hypothesised in the original study to be involved in establishing positional memory in zebrafish.

To characterise important biological processes of the DE transcripts, a selection group consisting of the 87 transcripts was created and subjected to gene ontology analysis using Goatools. Notably the GO term *oxidation-reduction process* (GO:0055114) was found to be significantly-enriched in the top-4 GO results for biological processes (p-value ≤ 0.05). Oxidation–reduction reactions are crucial for cell-growth and signalling and could play an important role in cellular regeneration [Balaban et al., 2005]. Among the genes that contributed to this GO term were *aldh1a2*, as well as the genes *pah* and *hgd* found in our results to be significantly changing in both the transcripts and protein levels. The differential expressions of *pah* and *hgd* at the protein level are consistent with existing literatures [Saxena et al., 2012], but from linked explorations, we observed that both *pah* and *hgd* were also DE at the transcript level. The results here could be investigated to gain further insights into the regulation mechanism of those genes.

Inspecting the linked Clustergrammer heatmaps of the DE transcripts and proteins (Supplementary Figure S2), clear block structures could be observed across the distally-enriched and proximally-enriched entities. These are the transcripts and proteins that could potentially contribute to patterned regeneration in zebrafish tissues. The clustering structure in the linked compounds are less clear, suggesting that the relationship between transcript and protein expression to metabolism during regeneration is a complex process. For more details, refer to Supplementary Figure S2

### Analysing Enriched Metabolic Pathways in Zebrafish

The original study [Rabinowitz et al., 2017] did not perform any pathway analysis. Using GraphOmics we investigated which metabolites and pathways contribute to positional memory and possibly regeneration. The Query Builder was used to filter for DE metabolites (as determined by *limma*) that are also linked to highly active pathways (as determined by PLAGE). A threshold of ≤ 0.05 was used on the p-values of both DE metabolites and pathways. This resulted in 45 DE metabolites spread across 57 DE pathways, listed in Supplementary Table S3. Among the significant pathways of interests are *Alanine metabolism* which makes sense as both alanine and glutamate were DE in the data. Consistent with the original study, Arginine is observed to be producing the largest DE amongst the significant compounds, alongside other compounds like glutamine and leucine. This is explained in the original study that as how they promote wound healing and encouraging cellular growth [Rabinowitz et al., 2017].

To obtain descriptive terms that characterise the overall biological processes of these metabolic pathways, we performed GO analysis on the 236 DE transcripts (p-values ≤ 0.01 and log fold changes at least ±0.5) that are linked to these DE compounds and pathways. The first two most significant biological process GO terms include *G protein-coupled receptor signaling pathway* (GO:0007186) and *signal transduction* (GO:0007165), showing that the activity level of signalling pathways are high. The findings here support the hypothesis in the original study on the influence of signalling pathways towards positional memory.

### Covid-19 Case Study

Understanding the Covid-19 disease on the molecular level through omics technologies could potentially offer new insights leading to the nature of the SARS-CoV-2 virus and the development of new treatments. Here we demonstrated how GraphOmics could be used to analyse and interactively explore the integrated results from a dual-omics (proteomics and metabolomics) study on the sera of Covid-19 patients [Shen et al., 2020].

### Data Loading and Pre-processing

The original study aimed to characterise the proteome and metabolome of a cohort of 28 severe Covid-19 patients in comparison to a cohort of 28 healthy patients. Processed proteins and metabolites data from the original study was retrieved. The protein data was provided in a format acceptable to GraphOmics (with rows identified by their UniProt ID) and could be readily uploaded. For metabolite data, each compound was identified by its chemical name in the original data. An automated script (available from our repository) was created to map from compound names to KEGG ID using the Bioservices library [Cokelaer et al., 2013]. Of the 905 names present in the original data, 220 could be matched based on matching by exact chemical names alone. This represented the majority of amino acids discussed in the original study, although it left out many lipid, steroid hormones and other chemicals that could not be easily mapped to KEGG and Reactome based on matching by exact chemical names alone.

Similar to the previously analysed Zebrafish data, DE analysis were performed on the Covid19 protein and metabolite data using *limma*, while PLAGE was used to analyse pathway activity levels on both omics types.

### Interactive Omics Exploration of the Covid19 Data

Once the initial data integration has been performed in GraphOmics, users could interactively explore the data to reveal biologically relevant hypotheses. Firstly to discover significantly changing entities, the Query Builder was used to filter for DE proteins (defined in the original study as having p-values ≤ 0.05 and log fold changes at least ±0.25), linked to pathways that are also significantly changing (p-values ≤ 0.05) based on the protein data were selected. This resulted in 139 proteins connected to 86 pathways, detailed in Supplementary Table S4.

Among the significant pathways in the results, two were related to the activation of the complement system, including *Terminal pathway of complement* (R-HSA-166665) and *Alternative complement activation* (R-HSA-173736). Note that while pathway analysis in the original study was performed using a completely different proprietary software [IPA, Krämer et al., 2014], our results are in agreement with how complement system was activated in the severe case in response to pathogens. Additionally the original study thoroughly discussed the high activity level of the *Platelet degranulation* (R-HSA-114608) pathway. This was also found to be significant in our results, and it could be explained by how platelets produced in the lung were activated in response to lung injury in the severe patients. All these significant pathways and their connections to DE entities can be browsed through GraphOmics.

We further illustrated how GraphOmics could identify other significant entities that are linked to those groups of DE proteins discovered above. Keeping the same filtering criteria, we selected the *Platelet degranulation* pathway from the Data Browser. This selected the DE proteins linked to that pathway and all their related entities. From the corresponding Clustergrammer view, two clusters of proteins that are either up-regulated or down-regulated in the severe-vs-healthy comparison could be observed (Supplementary Figure S5). The protein P02776 (for gene Platelet Factor 4, or PF4) was a member of the down-regulated cluster. The presence of PF4 in the down-regulated cluster was interesting because changes to PF4 was noted in the original study to be a prognosis marker in severe acute respiratory syndrome [Poon et al., 2012]. Its down-regulation in the severe group could support this hypothesis. Cropping this cluster in Clustergrammer resulted in a selection of the 17 member proteins and their connections to compounds, reactions and pathways in the Data Browser. This could be inspected to reveal additional relationships between entities. For example, the original study highlighted how serotonin level decreases with increasing severity of the disease as serotonin was transported to platelet for storage. The connection of serotonin to *Platelet degranulation* and to members of this cluster, and the down-regulation of serotonin could be interactively seen and explored from the Data Browser.

Finally we investigated the metabolomics data by filtering from the Query Builder for DE metabolites linked to DE pathways (p-values ≤ 0.05 for both). This resulted in 45 significant metabolites linked to 93 significant pathways. Examining the resulting metabolites, two clusters, one showing an up-regulation trend in the severe cohort, and one with down-regulation trend could be observed from Clustergrammer (Supplementary Figure S6). The first cluster contained kynurerine and NAD+. Its up-regulation was explained in the original study by the activation of kyunerine pathways in severe patients due to macrophage responses. The second down-regulated cluster contained many amino acids such as histidine, arginine, proline, and many others. Its down-regulation had been hypothesised in the original study to be due to damage to the liver from the disease.

## Discussion and Conclusions

In this work, we introduced GraphOmics, a Web application that could be used to explore and integrate biological data from the transcriptome, proteome and metabolome domains. Integration is achieved horizontally by mapping relevant biological entities to reactions and pathways from Reactome. Once mapping has been established, GraphOmics allows users to interrogate the data and interactively explore the connections between entities in the context of Reactome pathways.

To guide this exploration process, GraphOmics allows users to run several common global analyses, including differential expression and pathway activity analysis that prioritise DE entities in the data based on how they change across different experimental conditions. More interestingly, the connections between DE entities could also be explored and queried interactively within GraphOmics. The close integration between the Data Browser and interactive clustering and heatmaps in Clustergrammer means different views on the same data are synchronised to one another. This allows for integrated analysis where for instance, clustering results can be easily examined in the context of pathway activity levels.

Based on Reactome, GraphOmics supports as many species as Reactome offers. This is an advantage compared to other tools such as 3omics that supports human data only. Other tools like MetaboAnalyst and PaintOmics3 supports many species too, but they lack the easy inter-connectivity of results and close integration between multiple views in GraphOmics. As Reactome continues to grow, the knowledge base of GraphOmics also expands. Upgrading Reactome is as easy as pointing the GraphOmics server to an updated instance of the database.

As shown by the case studies on two complex multi-omics Zebrafish and Covid19 datasets, GraphOmics could be used to rapidly reveal interesting biological insights and potentially suggest relevant hypothesis. The first case study highlighted how users could use GraphOmics to find differentially expressed transcripts, proteins and metabolites involved in the caudal fin regeneration of zebrafish in agreement with the original study. Using the Covid19 data, we also demonstrated how users could use GraphOmics to reveal DE entities and pathways that were significantly changing in light of the disease. Here the results from GraphOmics were found consistent with findings in the original study. It is worth emphasising that throughout this entire process, omics data investigation and exploration in GraphOmics were performed interactively through the Web interface and did not require users to write manual R scripts for data analysis, as what were done in the original studies.

A weakness of GraphOmics is the requirement for entities to be identified and mapped to their IDs before they can be processed. While this requirement is more standard for transcript and protein data, it could be a challenge in metabolomics where a single compound could be associated to many chemical names and under different ID schemes. Additionally the uncertainty of peak annotations means a vast majority of metabolites in an untargeted study are not identified or could only be identified with a low level of confidence [Dunn et al., 2013, da Silva et al., 2015]. This is a weakness of nearly all tools that map metabolomics data to pathways. After the initial upload step, tools like MetaboAnalyst and PaintOmics3 display a screen for users to manually inspect, validate metabolite identities and delete duplicate annotations if they were present. This is a functionality that could be added to GraphOmics. Additionally, methods like Mummichog [Li et al., 2013] and PUMA [Hosseini et al., 2020] that combine metabolite annotation and pathway activity prediction steps together to increase confidence in the results could also be incorporated into GraphOmics.

Finally the integration approach in GraphOmics is currently restricted to only known entities and connections in Reactome. In the late integration approach adopted by GraphOmics, it is possible to miss the correlated entities that could have been discovered in an early integration scheme. To find the connections between unknown entities not present in the knowledge-base, methods such as correlation analysis, Bayesian analysis (e.g. MOFA [Argelaguet et al., 2018]), and other forms of latent factor analysis including clusterings of multi-omics data [Kirk et al., 2012, Lock and Dunson, 2013] could be employed. In the future we plan to extend GraphOmics to support factor-based analyses. This paves the way towards a platform that integrates data both horizontally (sharing common features) as well as vertically (sharing common samples) and presents the results in a truly integrated manner.

## Author’s contributions

JW and RD developed and tested the application. JW performed the analysis for the case studies. JW and RD wrote the manuscript. All authors read and approved the final version of the manuscript.

## Availability of data and materials

Datasets for the Zebrafish and Covid-19 case studies were obtained from the original studies and processed into a format suitable for GraphOmics analysis. Analysis results are available on GraphOmics and they can be accessed online. For the Zebrafish data, it is https://graphomics.glasgowcompbio.org/app/linker/explore_data/5, while for the Covid-19 data, it is https://graphomics.glasgowcompbio.org/app/linker/explore_data/6.

## References

Biswapriya B Misra, Carl Langefeld, Michael Olivier, and Laura A Cox. Integrated omics: tools, advances and future approaches. Journal of molecular endocrinology, 62(1):R21–R45, 2019.

Jason Lloyd-Price, Cesar Arze, Ashwin N Ananthakrishnan, Melanie Schirmer, Julian Avila-Pacheco, Tiffany W Poon, Elizabeth Andrews, Nadim J Ajami, Kevin S Bonham, Colin J Brislawn, et al. Multi-omics of the gut microbial ecosystem in inflammatory bowel diseases. Nature, 569(7758):655–662, 2019.

Suhas V Vasaikar, Peter Straub, Jing Wang, and Bing Zhang. Linkedomics: analyzing multi-omics data within and across 32 cancer types. Nucleic acids research, 46(D1):D956–D963, 2018.

Takoua Jendoubi. Approaches to integrating metabolomics and multi-omics data: A primer. Metabolites, 11(3):184, 2021.

Marinka Žitnik and Blaž Zupan. Data fusion by matrix factorization. IEEE transactions on pattern analysis and machine intelligence, 37(1):41–53, 2014.

Ricard Argelaguet, Britta Velten, Damien Arnol, Sascha Dietrich, Thorsten Zenz, John C Marioni, Florian Buettner, Wolfgang Huber, and Oliver Stegle. Multi-omics factor analysis—a framework for unsupervised integration of multi-omics data sets. Molecular systems biology, 14(6), 2018.

Zhiqiang Pang, Jasmine Chong, Guangyan Zhou, David Anderson de Lima Morais, Le Chang, Michel Barrette, Carol Gauthier, Pierre-Étienne Jacques, Shuzhao Li, and Jianguo Xia. Metaboanalyst 5.0: narrowing the gap between raw spectra and functional insights. Nucleic Acids Research, 2021.

Tien-Chueh Kuo, Tze-Feng Tian, and Yufeng Jane Tseng. 3omics: a web-based systems biology tool for analysis, integration and visualization of human transcriptomic, proteomic and metabolomic data. BMC systems biology, 7(1): 64, 2013.

Rafael Hernández-de Diego, Sonia Tarazona, Carlos Martínez-Mira, Leandro Balzano-Nogueira, Pedro Furió-Tarí, Georgios J Pappas Jr, and Ana Conesa. Paintomics 3: a web resource for the pathway analysis and visualization of multi-omics data. Nucleic acids research, 46(W1):W503–W509, 2018.

Ludovic Cottret, David Wildridge, Florence Vinson, Michael P Barrett, Hubert Charles, Marie-France Sagot, and Fabien Jourdan. Metexplore: a web server to link metabolomic experiments and genome-scale metabolic networks. Nucleic acids research, 38(suppl_2):W132–W137, 2010.

Hans-Jörg Schulz and Christophe Hurter. Grooming the hairball - how to tidy up network visualizations? In INFOVIS 2013, IEEE Information Visualization Conference, Atlanta, United States, October 2013. URL https://hal-enac.archives-ouvertes.fr/hal-00912739. Tutorial.

David Croft, Antonio Fabregat Mundo, Robin Haw, Marija Milacic, Joel Weiser, Guanming Wu, Michael Caudy, Phani Garapati, Marc Gillespie, Maulik R Kamdar, et al. The reactome pathway knowledgebase. Nucleic acids research, 42(D1):D472–D477, 2014.

Michael I Love, Wolfgang Huber, and Simon Anders. Moderated estimation of fold change and dispersion for rna-seq data with deseq2. Genome biology, 15(12):550, 2014.

Gordon K Smyth. Limma: linear models for microarray data. In Bioinformatics and computational biology solutions using R and Bioconductor, pages 397–420. Springer, 2005.

Andrew D Rouillard, Gregory W Gundersen, Nicolas F Fernandez, Zichen Wang, Caroline D Monteiro, Michael G McDermott, and Avi Ma’ayan. The harmonizome: a collection of processed datasets gathered to serve and mine knowledge about genes and proteins. Database, 2016, 2016.

DV Klopfenstein, Liangsheng Zhang, Brent S Pedersen, Fidel Ramírez, Alex Warwick Vesztrocy, Aurélien Naldi, Christopher J Mungall, Jeffrey M Yunes, Olga Botvinnik, Mark Weigel, et al. Goatools: A python library for gene ontology analyses. Scientific reports, 8(1):1–17, 2018.

Nicolas F Fernandez, Gregory W Gundersen, Adeeb Rahman, Mark L Grimes, Klarisa Rikova, Peter Hornbeck, and Avi Ma’ayan. Clustergrammer, a web-based heatmap visualization and analysis tool for high-dimensional biological data. Scientific data, 4:170151, 2017.

Karen McLuskey, Joe Wandy, Isabel Vincent, Justin JJ Van Der Hooft, Simon Rogers, Karl Burgess, and Rónán Daly. Ranking metabolite sets by their activity levels. Metabolites, 11(2):103, 2021.

Aravind Subramanian, Pablo Tamayo, Vamsi K Mootha, Sayan Mukherjee, Benjamin L Ebert, Michael A Gillette, Amanda Paulovich, Scott L Pomeroy, Todd R Golub, Eric S Lander, and Jill P Mesirov. Gene set enrichment analysis: a knowledge-based approach for interpreting genome-wide expression profiles. Proceedings of the National Academy of Sciences of the United States of America, 102(43):15545–15550, oct 2005.

John Tomfohr, Jun Lu, and Thomas B Kepler. Pathway level analysis of gene expression using singular value decomposition. BMC Bioinformatics, 6(1):225, 2005.

Adi L Tarca, Gaurav Bhatti, and Roberto Romero. A comparison of gene set analysis methods in terms of sensitivity, prioritization and specificity. PloS one, 8(11), 2013.

Antonio Fabregat, Konstantinos Sidiropoulos, Guilherme Viteri, Oscar Forner, Pablo Marin-Garcia, Vicente Arnau, Peter D’Eustachio, Lincoln Stein, and Henning Hermjakob. Reactome pathway analysis: a high-performance in-memory approach. BMC bioinformatics, 18(1):142, 2017.

Jeremy S Rabinowitz, Aaron M Robitaille, Yuliang Wang, Catherine A Ray, Ryan Thummel, Haiwei Gu, Danijel Djukovic, Daniel Raftery, Jason D Berndt, and Randall T Moon. Transcriptomic, proteomic, and metabolomic landscape of positional memory in the caudal fin of zebrafish. Proceedings of the National Academy of Sciences, 114 (5):E717–E726, 2017.

Robert S Balaban, Shino Nemoto, and Toren Finkel. Mitochondria, oxidants, and aging. cell, 120(4):483–495, 2005.

Sandeep Saxena, Sachin K Singh, Mula G Meena Lakshmi, Vuppalapaty Meghah, Bhawna Bhatti, Cherukuvada V Brahmendra Swamy, Curam S Sundaram, and Mohammed M Idris. Proteomic analysis of zebrafish caudal fin regeneration. Molecular & Cellular Proteomics, 11(6):M111–014118, 2012.

Bo Shen, Xiao Yi, Yaoting Sun, Xiaojie Bi, Juping Du, Chao Zhang, Sheng Quan, Fangfei Zhang, Rui Sun, Liujia Qian, et al. Proteomic and metabolomic characterization of covid-19 patient sera. 2020.

Thomas Cokelaer, Dennis Pultz, Lea M Harder, Jordi Serra-Musach, and Julio Saez-Rodriguez. Bioservices: a common python package to access biological web services programmatically. Bioinformatics, 29(24):3241–3242, 2013.

Andreas Krämer, Jeff Green, Jack Pollard Jr, and Stuart Tugendreich. Causal analysis approaches in ingenuity pathway analysis. Bioinformatics, 30(4):523–530, 2014.

Terence CW Poon, Ronald TK Pang, KC Allen Chan, Nelson LS Lee, Rossa WK Chiu, Yu-Kwan Tong, Stephen SC Chim, Sai M Ngai, Joseph JY Sung, and YM Dennis Lo. Proteomic analysis reveals platelet factor 4 and beta-thromboglobulin as prognostic markers in severe acute respiratory syndrome. Electrophoresis, 33(12):1894–1900, 2012.

Warwick B Dunn, Alexander Erban, Ralf JM Weber, Darren J Creek, Marie Brown, Rainer Breitling, Thomas Hankemeier, Royston Goodacre, Steffen Neumann, Joachim Kopka, et al. Mass appeal: metabolite identification in mass spectrometry-focused untargeted metabolomics. Metabolomics, 9(1):44–66, 2013.

Ricardo R da Silva, Pieter C Dorrestein, and Robert A Quinn. Illuminating the dark matter in metabolomics. Proceedings of the National Academy of Sciences, 112(41):12549–12550, 2015.

Shuzhao Li, Youngja Park, Sai Duraisingham, Frederick H Strobel, Nooruddin Khan, Quinlyn A Soltow, Dean P Jones, and Bali Pulendran. Predicting network activity from high throughput metabolomics. PLoS Computational Biology, 9(7):e1003123, 2013.

Ramtin Hosseini, Neda Hassanpour, Li-Ping Liu, and Soha Hassoun. Pathway-Activity Likelihood Analysis and Metabolite Annotation for Untargeted Metabolomics Using Probabilistic Modeling. Metabolites, 10(5):183, may 2020. ISSN 2218-1989. doi:10.3390/metabo10050183. URL https://www.mdpi.com/2218-1989/10/5/183.

Paul Kirk, Jim E Griffin, Richard S Savage, Zoubin Ghahramani, and David L Wild. Bayesian correlated clustering to integrate multiple datasets. Bioinformatics, 28(24):3290–3297, 2012.

Eric F Lock and David B Dunson. Bayesian consensus clustering. Bioinformatics, 29(20):2610–2616, 2013.

