## Supplementary Information for "GraphOmics: An Interactive Platform To Explore And Integrate Multi-Omics Data"

#### Supplementary Section S1: Input file format

To use GraphOmics, for each omics of interest, users have to prove its measurement data in Comma-separated Values (CSV) format. This takes the form of a matrix, where the first column contains the ID and further columns represent individual samples. For example, the intensity matrix takes the form of:

```
Identifier,sample1,sample2,sample3,sample4,FC_case1_vs_control,padj_case1_vs_control
ENSDARG00000000001,510,402,698,783,1.246,0.000284
ENSDARG00000000002,283,129,164,269,-0.7249,1.27e-06
ENSDARG00000000018,545,503,547,387,-0.0401,0.760679007
```

The following identifiers are accepted:

- Transcripts: Ensembl ID
- Proteins: UniProt ID
- Metabolites: KEGG or ChEBI ID

If statistical tests have been performed for the entities, the results can also be included as columns in the matrix. The column header has to be in the following format: 'FC_<xx>_vs_<yy>' for fold change information, and 'padj_<xx>_vs_<yy>' for p-values.

Finally a design matrix has to be specified to assign samples to factors (experimental conditions). This takes the form of another CSV file in this format:

```
sample,group
sample1,case1
sample2,case1
sample3,control
sample4,control
```

### Supplementary Figure S2: Clustergrammer Results for the Zebrafish Data

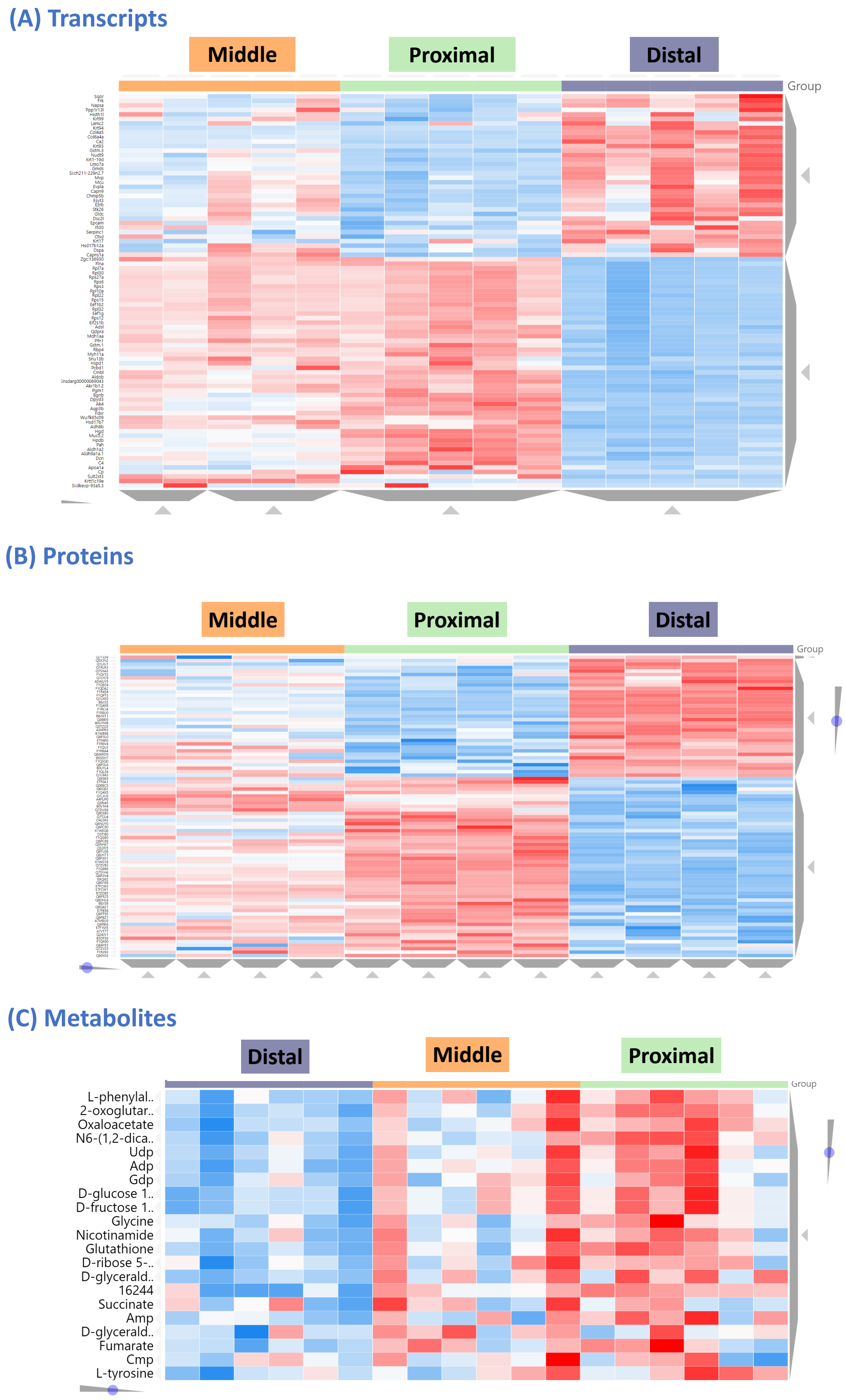


Clustergrammer results for selected transcripts, proteins and metabolites. The Query Builder was used to filter for transcripts and proteins with a threshold of 0.05 on the p-values, and at least ±0.5 on the log fold changes of the transcripts in the distal-vs-proximal comparison. The results were a selection of 87 transcripts and their corresponding proteins, as well as 21 compounds involved in reactions catalysed by those proteins.

#### Supplementary Table S3: Differentially Expressed Pathways from the Zebrafish Metabolite Data

| Reactome ID | Pathways | p-values transcripts | p-values proteins | p-values metabolites |
|---|---|---|---|---|
| R-DRE-5223345 | Miscellaneous transport and binding events | 3.11E-03 | - | 1.91E-03 |
| R-DRE-8964540 | Alanine metabolism | 9.82E-01 | - | 2.81E-03 |
| R-DRE-8963684 | Tyrosine catabolism | 7.46E-03 | 1.07E-02 | 3.50E-03 |
| R-DRE-352230 | Amino acid transport across the plasma membrane | 1.79E-03 | - | 3.80E-03 |
| R-DRE-420499 | Class c/3 (metabotropic glutamate/pheromone recep… | 2.10E-02 | - | 3.93E-03 |
| R-DRE-428559 | Proton-coupled neutral amino acid transporters | - | - | 4.28E-03 |
| R-DRE-71240 | Tryptophan catabolism | 1.92E-02 | - | 4.46E-03 |
| R-DRE-442660 | Na+/cl- dependent neurotransmitter transporters | 9.74E-04 | - | 4.59E-03 |
| R-DRE-8964539 | Glutamate and glutamine metabolism | 3.97E-02 | 1.00E+00 | 5.76E-03 |
| R-DRE-70921 | Histidine catabolism | 5.84E-03 | - | 5.93E-03 |
| R-DRE-8963693 | Aspartate and asparagine metabolism | 3.19E-02 | 1.00E+00 | 6.58E-03 |
| R-DRE-70895 | Branched-chain amino acid catabolism | 7.04E-02 | 1.00E+00 | 6.80E-03 |
| R-DRE-210455 | Astrocytic glutamate-glutamine uptake and metabol… | 1.60E-01 | - | 7.10E-03 |
| R-DRE-446210 | Synthesis of udp-n-acetyl-glucosamine | 7.19E-01 | 1.00E+00 | 7.45E-03 |
| R-DRE-156584 | Cytosolic sulfonation of small molecules | 7.10E-03 | 1.00E+00 | 8.11E-03 |
| R-DRE-561048 | Organic anion transport | 8.36E-01 | - | 8.48E-03 |
| R-DRE-5423646 | Aflatoxin activation and detoxification | 7.59E-03 | - | 9.32E-03 |
| R-DRE-174403 | Glutathione synthesis and recycling | 6.94E-03 | - | 9.82E-03 |
| R-DRE-2142691 | Synthesis of leukotrienes (lt) and eoxins (ex) | 2.26E-03 | - | 9.82E-03 |
| R-DRE-5683826 | Surfactant metabolism | 9.41E-03 | 3.49E-01 | 1.04E-02 |
| R-DRE-71288 | Creatine metabolism | 2.60E-02 | - | 1.05E-02 |
| R-DRE-549127 | Organic cation transport | 9.83E-04 | - | 1.07E-02 |
| R-DRE-196807 | Nicotinate metabolism | 6.22E-02 | - | 1.15E-02 |
| R-DRE-196757 | Metabolism of folate and pterines | 6.81E-02 | 4.71E-01 | 1.23E-02 |
| R-DRE-416476 | G alpha (q) signalling events | 5.72E-04 | 1.88E-01 | 1.36E-02 |
| R-DRE-5628897 | Tp53 regulates metabolic genes | 7.47E-01 | 9.79E-01 | 1.36E-02 |
| R-DRE-71064 | Lysine catabolism | 8.81E-02 | 1.00E+00 | 1.41E-02 |
| R-DRE-389661 | Glyoxylate metabolism and glycine degradation | 9.38E-02 | 7.85E-01 | 1.59E-02 |
| R-DRE-70635 | Urea cycle | 3.48E-01 | - | 1.61E-02 |
| R-DRE-9639288 | Amino acids regulate mtorc1 | 1.00E+00 | 1.00E+00 | 1.68E-02 |
| R-DRE-500753 | Pyrimidine biosynthesis | 1.92E-01 | - | 1.72E-02 |
| R-DRE-163754 | Insulin effects increased synthesis of xylulose-5… | - | - | 1.84E-02 |
| R-DRE-70263 | Gluconeogenesis | 2.24E-02 | 8.39E-02 | 1.85E-02 |
| R-DRE-71336 | Pentose phosphate pathway | 1.32E-01 | 4.75E-01 | 1.86E-02 |
| R-DRE-70350 | Fructose catabolism | 5.16E-03 | - | 1.90E-02 |
| R-DRE-196783 | Coenzyme a biosynthesis | 2.63E-02 | - | 1.98E-02 |
| R-DRE-70171 | Glycolysis | 2.97E-02 | 2.15E-01 | 2.13E-02 |
| R-DRE-418594 | G alpha (i) signalling events | 5.50E-04 | 1.41E-01 | 2.23E-02 |
| R-DRE-8964208 | Phenylalanine metabolism | 2.95E-03 | 7.06E-03 | 2.23E-02 |
| R-DRE-70370 | Galactose catabolism | 1.00E+00 | - | 2.30E-02 |
| R-DRE-156581 | Methylation | 5.50E-03 | 1.00E+00 | 2.30E-02 |
| R-DRE-428643 | Organic anion transporters | 2.01E-01 | - | 2.40E-02 |
| R-DRE-159418 | Recycling of bile acids and salts | 1.45E-02 | - | 2.42E-02 |
| R-DRE-193368 | Synthesis of bile acids and bile salts via 7alpha… | 2.96E-02 | - | 2.42E-02 |
| R-DRE-70221 | Glycogen breakdown (glycogenolysis) | 2.41E-03 | - | 2.49E-02 |
| R-DRE-916853 | Degradation of gaba | - | - | 2.58E-02 |
| R-DRE-3928662 | Ephb-mediated forward signaling | 5.30E-01 | 9.99E-01 | 2.64E-02 |
| R-DRE-165159 | Mtor signalling | 7.68E-01 | - | 2.78E-02 |
| R-DRE-5673001 | Raf/map kinase cascade | 1.21E-03 | 1.00E+00 | 2.97E-02 |
| R-DRE-418555 | G alpha (s) signalling events | 2.93E-05 | 1.96E-01 | 3.01E-02 |
| R-DRE-71403 | Citric acid cycle (tca cycle) | 5.60E-01 | 5.98E-02 | 3.10E-02 |
| R-DRE-389599 | Alpha-oxidation of phytanate | 1.68E-01 | - | 4.24E-02 |
| R-DRE-1660662 | Glycosphingolipid metabolism | 1.70E-02 | 1.00E+00 | 4.26E-02 |
| R-DRE-71262 | Carnitine synthesis | 1.67E-02 | - | 4.36E-02 |
| R-DRE-5625886 | Activated pkn1 stimulates transcription of ar (an… | 9.93E-01 | 1.00E+00 | 4.75E-02 |

| | | | | |
|---|---|---|---|---|
| R-DRE-2299718 | Condensation of prophase chromosomes | 9.93E-01 | 1.81E-01 | 4.75E-02 |
| R-DRE-2559580 | Oxidative stress induced senescence | 1.48E-02 | 5.47E-01 | 4.75E-02 |

The 57 DE pathways that are linked to DE metabolites in the Zebrafish data. *limma* was applied to assess the significance of DE metabolites, while PLAGE was applied on the metabolite data to prioritise pathway activities in the distal-vs-proximal comparison. For each pathway, a minimum hits of 3 entities was required for PLAGE results to be considered (results with less than 3 hits are represented by a '-' in the table). The Query Builder in GraphOmics was used to filter for DE metabolites that belonged to highly active pathways (based on the metabolite data). A threshold of $\leq 0.05$ was used for the p-values. This resulted in 45 compounds across 57 pathways listed above. PLAGE results are shown in the *p-values metabolites* columns and used to sort the table. Additionally the results of running PLAGE on the transcripts and proteins data are also shown in the table under the *p-values transcripts* and *p-values proteins* columns respectively.

#### Supplementary Table S4: Differentially Expressed Pathways from the Covid-19 Protein Data

| Reactome ID | Pathways | p-values proteins | p-values reactome |
|---|---|---|---|
| R-HSA-354192 | Integrin signaling | 0.00E+00 | 2.11E-04 |
| R-HSA-8936459 | Runx1 regulates genes involved in megakaryocyte d... | 0.00E+00 | 1.89E-03 |
| R-HSA-372708 | P130cas linkage to mapk signaling for integrins | 0.00E+00 | 7.69E-06 |
| R-HSA-354194 | Grb2:sos provides linkage to mapk signaling for i... | 0.00E+00 | 4.34E-06 |
| R-HSA-114608 | Platelet degranulation | 0.00E+00 | 0.00E+00 |
| R-HSA-8964058 | Hdl remodeling | 0.00E+00 | 1.64E-04 |
| R-HSA-2022857 | Keratan sulfate degradation | 0.00E+00 | 1.43E-01 |
| R-HSA-3656225 | Defective chst6 causes mcdc1 | 0.00E+00 | 6.25E-02 |
| R-HSA-3656243 | Defective st3gal3 causes mct12 and eiee15 | 0.00E+00 | 6.25E-02 |
| R-HSA-977225 | Amyloid fiber formation | 0.00E+00 | 3.41E-05 |
| R-HSA-166016 | Toll like receptor 4 (tlr4) cascade | 0.00E+00 | 9.61E-02 |
| R-HSA-212300 | Prc2 methylates histones and dna | 0.00E+00 | 5.79E-02 |
| R-HSA-912446 | Meiotic recombination | 0.00E+00 | 9.56E-02 |
| R-HSA-2299718 | Condensation of prophase chromosomes | 0.00E+00 | 8.55E-02 |
| R-HSA-5617472 | Activation of anterior hox genes in hindbrain dev... | 0.00E+00 | 3.66E-01 |
| R-HSA-427359 | Sirt1 negatively regulates rrna expression | 0.00E+00 | 1.50E-02 |
| R-HSA-73728 | Rna polymerase i promoter opening | 0.00E+00 | 1.05E-01 |
| R-HSA-5334118 | Dna methylation | 0.00E+00 | 3.40E-02 |
| R-HSA-9616222 | Transcriptional regulation of granulopoiesis | 0.00E+00 | 1.54E-01 |
| R-HSA-427413 | Norc negatively regulates rrna expression | 0.00E+00 | 1.43E-01 |
| R-HSA-5625886 | Activated pkn1 stimulates transcription of ar (an... | 0.00E+00 | 1.43E-01 |
| R-HSA-2559582 | Senescence-associated secretory phenotype (sasp) | 0.00E+00 | 1.54E-01 |
| R-HSA-73772 | Rna polymerase i promoter escape | 0.00E+00 | 1.54E-01 |
| R-HSA-201722 | Formation of the beta-catenin:tcf transactivating... | 0.00E+00 | 1.12E-01 |
| R-HSA-1912408 | Pre-notch transcription and translation | 0.00E+00 | 1.54E-01 |
| R-HSA-427389 | Ercc6 (csb) and ehmt2 (g9a) positively regulate r... | 0.00E+00 | 6.46E-02 |
| R-HSA-5578749 | Transcriptional regulation by small rnas | 0.00E+00 | 1.92E-01 |
| R-HSA-9609690 | Hcmv early events | 0.00E+00 | 7.64E-01 |
| R-HSA-445989 | Tak1 activates nfkb by phosphorylation and activa... | 0.00E+00 | 5.16E-01 |
| R-HSA-933542 | Traf6 mediated nf-kb activation | 0.00E+00 | 4.17E-01 |
| R-HSA-879415 | Advanced glycosylation endproduct receptor signal... | 0.00E+00 | 2.57E-01 |
| R-HSA-3214815 | Hdacs deacetylate histones | 4.18E-19 | 1.54E-01 |
| R-HSA-6802946 | Signaling by moderate kinase activity braf mutants | 3.85E-18 | 1.29E-03 |
| R-HSA-5674135 | Map2k and mapk activation | 3.85E-18 | 7.50E-04 |
| R-HSA-6802952 | Signaling by braf and raf fusions | 3.85E-18 | 5.93E-03 |
| R-HSA-6802948 | Signaling by high-kinase activity braf mutants | 3.85E-18 | 4.27E-04 |
| R-HSA-6802955 | Paradoxical activation of raf signaling by kinase... | 3.85E-18 | 1.29E-03 |
| R-HSA-9649948 | Signaling downstream of ras mutants | 3.85E-18 | 1.29E-03 |
| R-HSA-418594 | G alpha (i) signalling events | 2.16E-17 | 2.60E-01 |
| R-HSA-9018519 | Estrogen-dependent gene expression | 2.88E-17 | 3.22E-01 |
| R-HSA-917937 | Iron uptake and transport | 3.00E-10 | 8.55E-02 |
| R-HSA-1221632 | Meiotic synapsis | 3.00E-10 | 3.20E-01 |
| R-HSA-3214847 | Hats acetylate histones | 5.00E-10 | 3.36E-01 |
| R-HSA-2559580 | Oxidative stress induced senescence | 8.00E-10 | 1.64E-01 |
| R-HSA-5250924 | B-wich complex positively regulates rrna expressi... | 1.20E-09 | 1.54E-01 |
| R-HSA-975634 | Retinoid metabolism and transport | 1.23E-08 | 1.92E-03 |
| R-HSA-3000178 | Ecm proteoglycans | 1.59E-08 | 1.92E-03 |
| R-HSA-8957275 | Post-translational protein phosphorylation | 2.23E-07 | 0.00E+00 |
| R-HSA-5694530 | Cargo concentration in the er | 2.25E-07 | 7.78E-03 |
| R-HSA-174577 | Activation of c3 and c5 | 4.45E-07 | 7.80E-09 |
| R-HSA-204005 | Copii-mediated vesicle transport | 6.07E-07 | 7.63E-02 |
| R-HSA-6785807 | Interleukin-4 and interleukin-13 signaling | 9.67E-07 | 1.54E-01 |
| R-HSA-381426 | Regulation of insulin-like growth factor (igf) tr... | 3.80E-06 | 0.00E+00 |
| R-HSA-166665 | Terminal pathway of complement | 9.60E-06 | 3.55E-04 |
| R-HSA-8963888 | Chylomicron assembly | 1.74E-05 | 2.13E-03 |

| | | | |
|---|---|---|---|
| R-HSA-2160916 | Hyaluronan uptake and degradation | 1.88E-05 | 1.35E-01 |
| R-HSA-75892 | Platelet adhesion to exposed collagen | 1.95E-05 | 3.28E-03 |
| R-HSA-8939236 | Runx1 regulates transcription of genes involved i... | 2.51E-05 | 2.82E-02 |
| R-HSA-446353 | Cell-extracellular matrix interactions | 3.41E-05 | 1.43E-01 |
| R-HSA-8963901 | Chylomicron remodeling | 3.84E-05 | 3.91E-04 |
| R-HSA-3656244 | Defective b4galt1 causes b4galt1-cdg (cdg-2d) | 4.07E-05 | 7.78E-03 |
| R-HSA-430116 | Gp1b-ix-v activation signalling | 9.29E-05 | 1.36E-03 |
| R-HSA-5686938 | Regulation of tlr by endogenous ligand | 2.34E-04 | 1.50E-04 |
| R-HSA-76009 | Platelet aggregation (plug formation) | 2.48E-04 | 2.60E-05 |
| R-HSA-3000480 | Scavenging by class a receptors | 3.45E-04 | 1.43E-01 |
| R-HSA-6798695 | Neutrophil degranulation | 8.39E-04 | 0.00E+00 |
| R-HSA-173736 | Alternative complement activation | 1.15E-03 | 2.74E-03 |
| R-HSA-5689880 | Ub-specific processing proteases | 1.72E-03 | 3.38E-01 |
| R-HSA-189483 | Heme degradation | 1.77E-03 | 8.76E-03 |
| R-HSA-1989781 | Ppara activates gene expression | 2.37E-03 | 2.28E-01 |
| R-HSA-2453902 | The canonical retinoid cycle in rods (twilight vi... | 4.85E-03 | 1.54E-01 |
| R-HSA-445355 | Smooth muscle contraction | 6.85E-03 | 2.32E-01 |
| R-HSA-8964041 | Ldl remodeling | 7.19E-03 | 6.00E-02 |
| R-HSA-3000170 | Syndecan interactions | 8.96E-03 | 1.90E-02 |
| R-HSA-186797 | Signaling by pdgf | 1.03E-02 | 1.54E-01 |
| R-HSA-8963889 | Assembly of active lpl and lipc lipase complexes | 1.23E-02 | 8.55E-02 |
| R-HSA-4420097 | Vegfa-vegfr2 pathway | 1.55E-02 | 1.10E-01 |
| R-HSA-1236977 | Endosomal/vacuolar pathway | 1.66E-02 | 4.51E-01 |
| R-HSA-9033241 | Peroxisomal protein import | 1.96E-02 | 7.13E-01 |
| R-HSA-3000471 | Scavenging by class b receptors | 2.08E-02 | 1.43E-01 |
| R-HSA-1650814 | Collagen biosynthesis and modifying enzymes | 2.09E-02 | 4.13E-01 |
| R-HSA-416476 | G alpha (q) signalling events | 2.75E-02 | 2.76E-01 |
| R-HSA-1474228 | Degradation of the extracellular matrix | 3.31E-02 | 8.74E-02 |
| R-HSA-202424 | Downstream tcr signaling | 3.60E-02 | 2.01E-01 |
| R-HSA-381038 | Xbp1(s) activates chaperone genes | 4.12E-02 | 8.30E-01 |
| R-HSA-71336 | Pentose phosphate pathway | 4.48E-02 | 1.43E-01 |

86 DE pathways that are linked to DE proteins in the Covid-19 data. *limma* was applied to assess protein significance, while PLAGE was applied on the protein data to prioritise pathway activities in the severe-vs-healthy comparison. For each pathway, a minimum hits of 3 entities was required for PLAGE results to be considered (results with less than 3 hits are represented by a '-' in the table). From the Query Builder in GraphOmics, significantly changing DE proteins (defined in the original study as having p-values $\leq 0.05$ and log fold changes at least $\pm 0.25$), linked to pathways that are also significantly changing (p-values $\leq 0.05$) were selected. This resulted in 139 proteins connected to 86 pathways listed above. PLAGE results are shown in the *p-values proteins* columns and used to sort the table. Additionally the results of running Reactome Analysis Service that considers both proteins and metabolites data for over-representation analysis are also shown under the *p-values reactome* column.

### Supplementary Figure S5: Differentially Expressed Proteins Linked to Platelet Degranulation Pathway in the Covid-19 Data

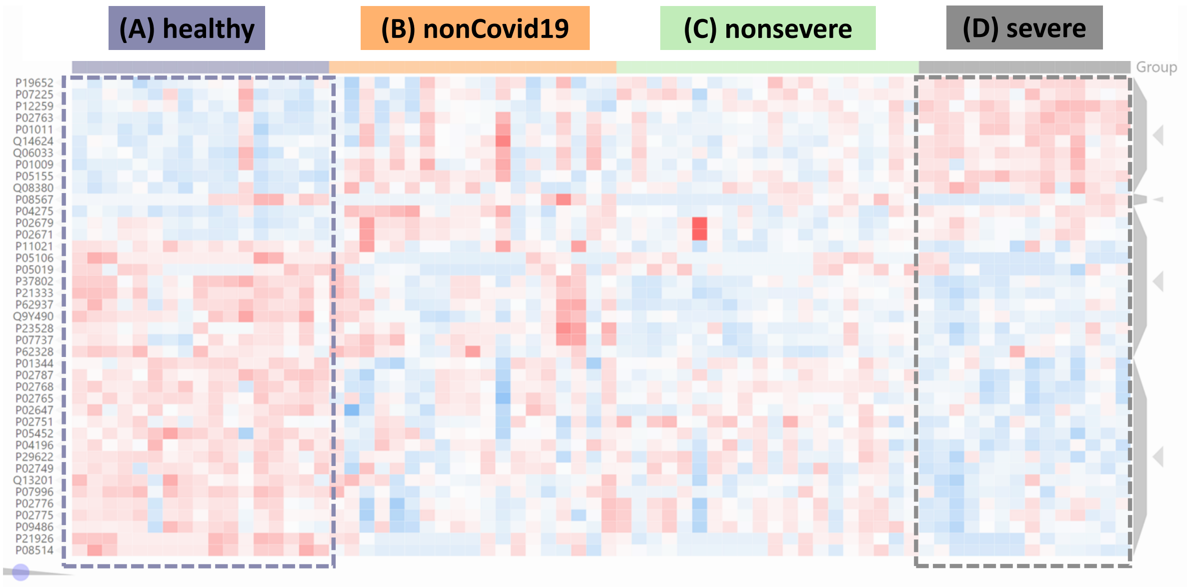


Clustergrammer results for the DE proteins linked to *Platelet Degranulation* pathway. From the Query Builder, significantly changing DE proteins (defined in the original study as having p-values $\leq 0.05$ and log fold changes at least $\pm 0.25$), linked to pathways that are also significantly changing (p-values $\leq 0.05$) were selected. From this, we filtered for proteins involved in *Platelet Degranulation* pathway from the Data Browser. Two clusters of up-/down-regulated proteins could be observed in the severe-vs-healthy comparison of DE proteins linked to *Platelet Degranulation*.

#### Supplementary Figure S6: Differentially Expressed Metabolites Linked to Differentially Expressed Pathways in the Covid-19 Data

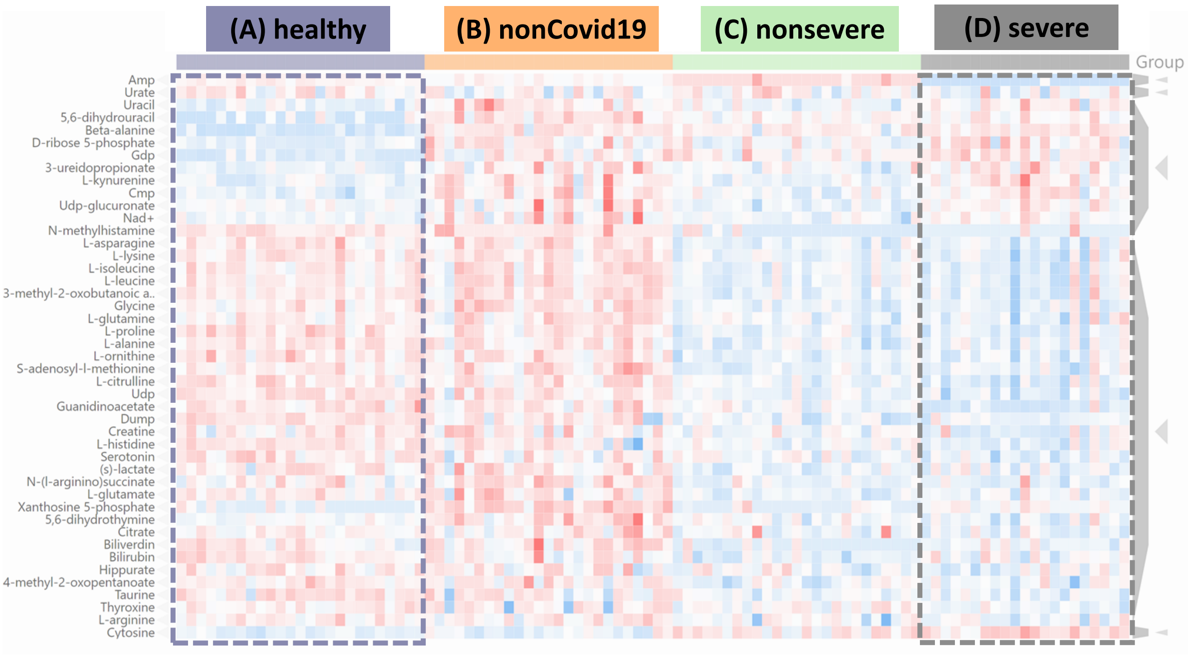


Clustergrammer results for the DE metabolites linked to DE pathways. From the Query Builder, we filtered for DE metabolites linked to DE pathways (p-values $\leq 0.05$ for both). Two clusters, one showing an upregulation trend in the severe cohort, and one with downregulation trend could be observed from the Clustergrammer results.
